# Searching for structure in collective systems

**DOI:** 10.1101/362681

**Authors:** Colin R. Twomey, Andrew T. Hartnett, Matthew M. Grobis, Pawel Romanczuk

## Abstract

Collective systems such as fish schools, bird flocks, and neural networks are comprised of many mutually-influencing individuals, often without long-term leaders, well-defined hierarchies, or persistent relationships. The remarkably organized group-level behaviors readily observable in these systems contrast with the ad hoc, often difficult to observe, and complex interactions among their constituents. While these complex individual-level dynamics are ultimately the drivers of group-level coordination, they do not necessarily offer the most parsimonious description of a group’s macroscopic properties. Rather, the factors underlying group organization may be better described at some intermediate, mesoscopic scale. We introduce a novel method from information-theoretic first principles to find a compressed description of a system based on the actions and mutual dependencies of its constituents, thus revealing the natural structure of the collective. We emphasize that this method is computationally tractable and requires neither pairwise nor Gaussian assumptions about individual interactions.

## 1 Introduction

Collective behavior is an emergent property of the actions and interactions of a system’s constituents. Typically, these individual actions are readily observable, while interactions are hidden: they can only be inferred indirectly, and in many cases only with great difficulty. A growing body of research on collective systems is devoted to exactly this problem (see e.g. Ballerini et al., 2008; Lukeman et al., 2010; Nagy et al., 2010; Katz et al., 2011; Herbert-Read et al., 2011; Bialek et al., 2012; Strandburg-Peshkin et al., 2013; Rosenthal et al., 2015; Harpaz et al., 2017; Torney et al., 2018). In general, methods for inferring interactions fall into one of two categories. Many make strong, system-specific simplifying assumptions about the nature of the interactions, e.g. requiring linearity and/or pairwise interaction topologies, thus limiting their widespread applicability. Other solutions relax these constraints but at the cost of tractability as systems scale in size to more than a few individuals. Even once accomplished, the problem of inferring all dependencies between all elements of a system at every moment is only the first step in the analysis of collective behavior. The overarching goal is to then understand how the dependencies between elements determine group-level coordination.

We will call this focus on characterizing the moment-to-moment interactions of a group the ‘bottom-up’ approach to quantifying collectivity. At the other end of the scale, the ‘top-down’ approach is to simply measure one or more bulk properties of the system, such as average alignment (when such a property makes sense; e.g. for locusts or fish in Buhl et al., 2006; Tunstrøm et al., 2013, respectively). However, for nest-site selection in honeybees (Seeley & Visscher, 2004), bridge formation (Reid et al., 2015) or foraging decisions (Greene & Gordon, 2007) in ants, social conflict policing in Macaques (Flack et al., 2006), quorum sensing in bacteria (Papenfort & Bassler, 2016), or neuronal avalanches in slices of neocortex (Beggs & Plenz, 2003), average alignment would not be the most meaningful aggregate measure of collectivity. In general, the choice of what bulk property to measure, and even what bulk properties may be sensible to measure, is system dependent.

The top-down and bottom-up views of collectivity are not mutually exclusive. On the contrary, ideally they are complementary, and it may be necessary to employ either or both depending on the system and the question asked. Here, we introduce a third approach to the problem of quantifying collectivity with the aim of unifying these two views, while addressing some of their practical and fundamental issues. First, building from information-theoretic first principles, we introduce a measure of aggregate collectivity based directly on the observable actions of a system’s individual elements. This measure quantifies the relative degree of statistical dependence shared by a set of elements and in principle is valid for any system of any size. The degree of macroscopic collectivity and its variation over time can thus be productively quantified and compared across systems. Second, we show that this measure allows us to find a natural decomposition of a system into simpler components. This decomposition provides a mesoscale description of a system that may offer a simpler basis on which to make inferences about the causal system-wide interactions that underly group-level organization. Finally, in addition to a rigorous theoretical foundation, we show that this approach is readily applicable to both observed and simulated experimental data in practice.

### 1.1 Redundancy

Let *S* = {1,2, …, *n*} be the indices of a set of random variables, {*X_i_*}*_i_*_∈*S*_, which in general may be neither identically distributed nor independent. In the context of a fish school or a bird flock, this could be the set of all the velocity vectors of the individuals in the group; for neurons, this could be the state of each neuron (firing or silent). In general, it could be any heterogeneous assemblage of the microscopic observables of a system. If we were asked to faithfully record the current state of the whole group, one strategy would be to simply write down a description of each element separately. One of the foundational results from information theory is that no lossless description of a random variable can be shorter on average than the tight lower bound given by its entropy (Shannon, 1948). Thus a description of the system given by recording every element separately would require on average a minimum of ∑*_i_*_∈*S*_ *H*(*X_i_*) bits, where *H*(*X_i_*) is the entropy of *X_i_*.

Alternatively, another strategy would be to instead write down a shared (or ‘joint’) description of all elements at once. A joint description can capitalize on the dependencies among a set of variables to reduce the overall description length needed. For example, to characterize the state of both a lamp and the light switch that controls it, one could simply record the on/off state of one of the two components. Knowing the state of either the switch or the lamp automatically tells us the state of the other, under perfect operating conditions. For less than perfect operating conditions it will be necessary to include additional information about the state of the other component, but only as frequently as the light switch fails to determine the state of the lamp. In either case, the joint entropy of the lamp and the light switch together determines the lower bound on the lossless joint description of the system. Thus the smallest lossless joint description requires *H*({*X_i_*}*_i_*_∈*S*_) bits on average, where we are guaranteed that *H*({*X_i_*}*_i_*_∈*S*_) ≤ ∑*_i_*_∈*S*_ *H*(*X_i_*).

In fact, the only way in which the joint description is as costly as the sum of the individual (or ‘marginal’) descriptions is if all *X_i_*’s are independent. The difference between the marginal and joint descriptions, given by

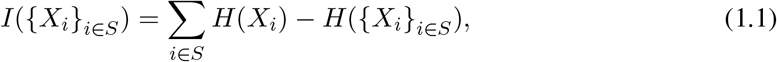

gives us a natural measure of how much we reduce the fundamental representation cost by using a joint, rather than a marginal, description. Another way to think about Eq. 1.1 is as a measure of redundancy: the amount of information that is made redundant (unnecessary) when describing {*X_i_*}*_i_*_∈*S*_ as a whole rather than by parts. A similar interpretation can be found in Watanabe (1960)’s original investigation of Eq. 1.1 as a general measure of multivariate correlation.^1^

Notably, redundancy in the absolute sense given by Eq. 1.1 scales in magnitude with the size of the system. For example, if we take *n* identical copies^2^ of the same random variable, *X*, then we have *I*({*X_i_*}*_i_*_∈*S*_) ∝ *H*(*X*) with the constant of proportionality equal to *n* − 1. If *H*(*X*) *>* 0, then in the limit as *n* → ∞, *I*({*X_i_*}*_i_*_∈*S*_) → ∞. To compare redundancies between systems or subsystems of different sizes, it can be useful to instead consider the relative redundancy, i.e.

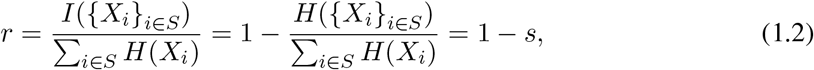

where *s* is then the proportion of non-redundant, or incompressible, information in the set. Taking the same example as before, for *n* identical copies of *X*, as *n → ∞*, *r →* 1, and *s* → 0 (see Fig. 1). At the other extreme, if instead of *n* identical copies we have *n* mutually independent *X_i_*’s, then as *n → ∞*, *r* = 0 and *s* = 1. In general, 0 ≤ *r* < 1 for any finite set of random variables with at least one variable having non-zero entropy, and, correspondingly, 0 *< s* ≤ 1.

**Figure 1:**
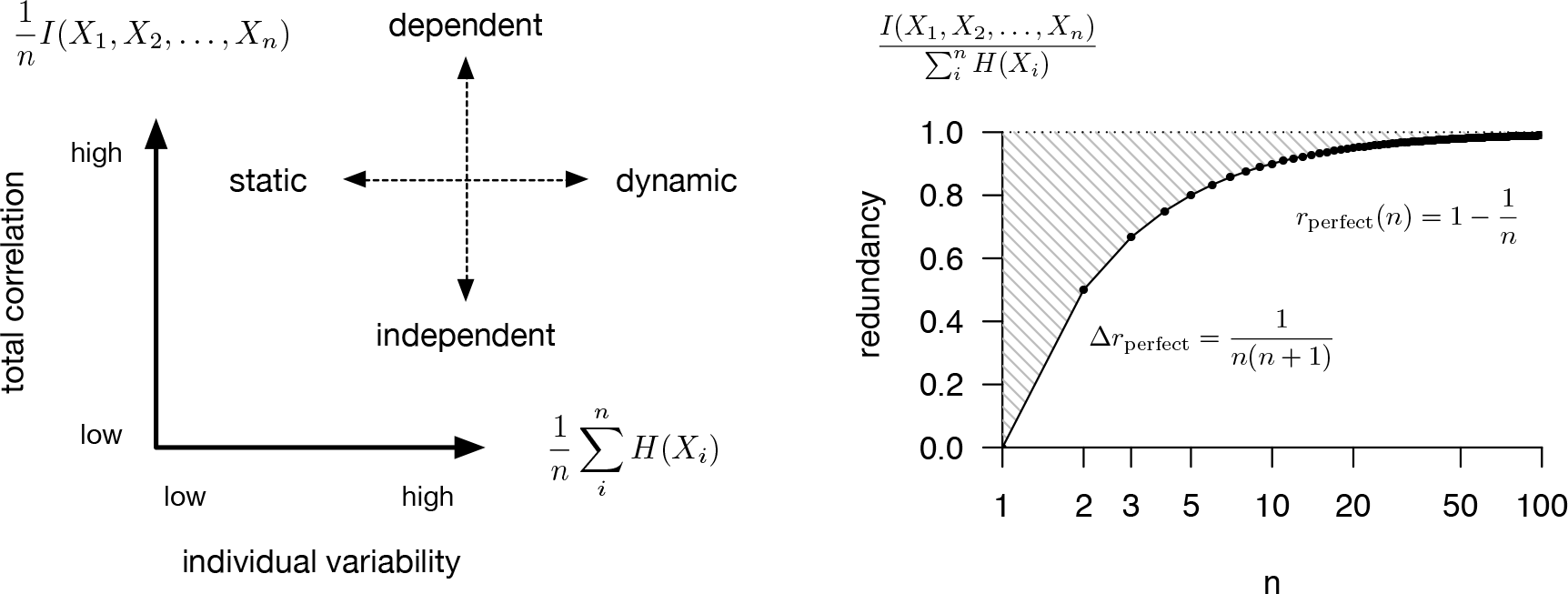
(*Left*) Schematic description of a system, {*X_i_*}*_i_*_∈*S*_, by its average total correlation (y-axis), measuring dependence, and the average marginal entropy of its elements (x-axis). (*Right*) Feasible (white) and infeasible (shaded) redundancies for systems of a given size, *n*. The upper bound is given by a system in which every element is perfectly dependent on every other element (so knowing the state of one element is as good as knowing the state of every element in the system). The lower bound is zero, which occurs when every element is independent of every other element.

## 2 Method

While relative redundancy, or equivalently incompressibility, can be used to compare the degree of collectivity exhibited by very different systems, it can also be used to characterize the dependency structure within a given system. Writing the relative redundancy as a function of a subset of the system, *A* ⊆ *S*, we have

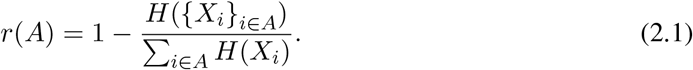

What divisions of a system maximize the relative redundancy of each subset? Can the subdivisions of a system achieve a higher average relative redundancy than the system as a whole?

To begin making these questions concrete, let *Ŝ* be a set of indices for a collection of subsets of *S*, which we will refer to as the *components* of system *S*. That is, let *Ŝ* = {1, 2, …, *m*}, where typically^3^ *m ≤ n*, and introduce a probabilistic assignment *p*(*j|i*), ∀(*i, j*) ∈ (*S,Ŝ*).^4^ Then the expected quality of an assignment to a given component is

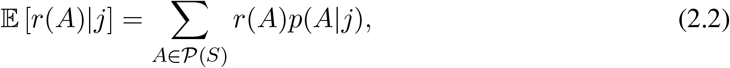

where 𝒫(*S*) is the power set (set of all subsets) of *S*, and

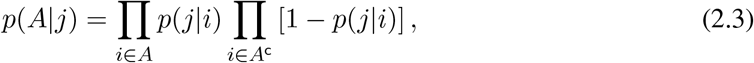

is the probability of subset *A* given the assignments of elements to component *j*, by a simple counting argument.^5^ Treating the quality of each component equally, the expected quality over all components is then

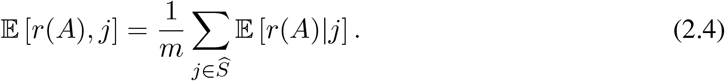

### 2.1 Rate-distortion theory

While this gives us a natural way to evaluate the quality of a given assignment, it does not immediately provide us with a way to find such an assignment. Instead, we draw inspiration from the information-theoretic treatment of compression given by rate-distortion theory (see Shannon, 1959; Cover & Thomas, 2006). Classical rate-distortion theory addresses the following problem: given a source (random variable) *X*, a measure of distortion, *d*, and an allowable level of average distortion *D*, determine the minimum rate necessary for a compressed description of *X* that introduces an average distortion no more than *D*. I.e.,

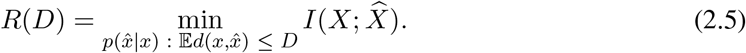

In this case, the rate measures the information, *I*(*X*; *X̂*), that the compressed representation, *X̂*, needs to keep about the source, *X*, where

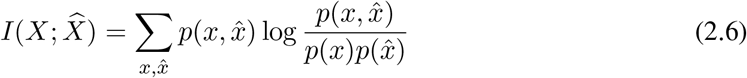

is the mutual information between *X* and *X̂*. The lower the rate, the better the compression, but (depending on the source and the distortion measure) the higher the average distortion introduced. Surprisingly, not only can the rate-distortion curve be characterized numerically in general, the minimal compressed representation of *X* can be found via a simple, iterative, alternating minimization algorithm (Blahut, 1972; Arimoto, 1972).

### 2.2 Redundancy compression

Though there are important differences from rate-distortion theory (discussed in Appendix A), we can similarly frame the problem of finding structure based on redundancy as a compression problem. Here, we wish to find the assignment of elements of *S* to components of *Ŝ* that achieves an average redundancy no less than *r*\*, and otherwise preserves as little about the original identities of the elements as possible. I.e.,

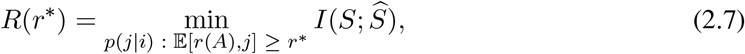

where *p*(*j|i*) is further required to be non-negative and sum to one. This is not a standard rate-distortion problem,^6^ but we can use much of the same ideas developed by Blahut (1972) and Arimoto (1972) in their original numerical algorithms for the channel capacity and rate-distortion problems for deriving a practical solution. We give a brief account of this derivation here; see Appendix A for a complete account.

Introducing Lagrange multipliers to constrain the ∑*_i_*_∈*S*_ *p*(*j|i*) = 1 (non-negativity will be enforced by the form of the solution), the variational problem becomes

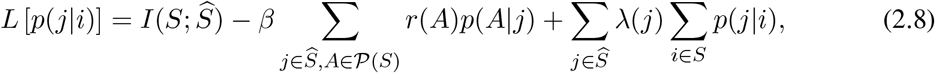

where *β*, the lagrange multiplier for the average redundancy constraint, absorbs the 1*/m* term. Taking the derivative with respect to a particular *j′* and *i′*, we have

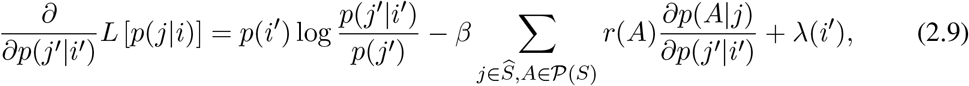

where

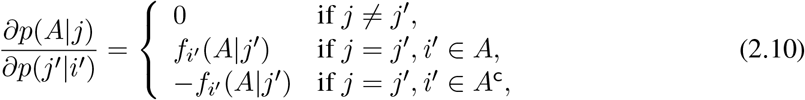

and

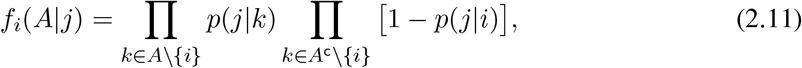

where *A* \ {*i*} is the relative complement of {*i*} with respect to *A*.

Then setting *∂L*/*∂p*(*j′|i′*) = 0 and splitting the sum over 𝒫(*S*) into terms with and without *i*′ ∈ *A*, we have

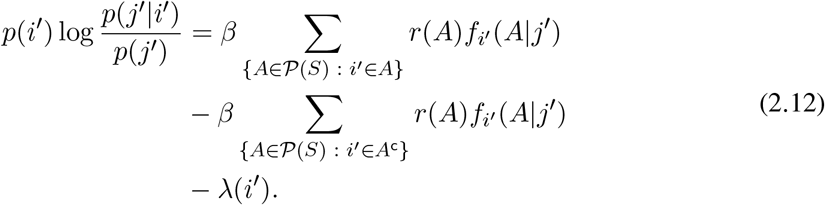

Let

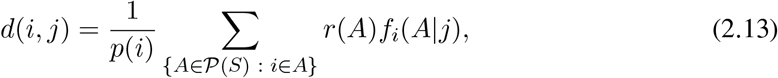

and define *d*_c_(*i, j*) to be identical except substituting *i* ∈ *A*^c^ for *i* ∈ *A*. Lastly, let Δ *d*(*i, j*) = *d*(*i, j*) *− d*_c_(*i, j*). Then, dividing through by *p*(*i′*) and substituting, we have,

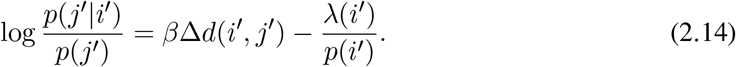

Finally, substituting log *µ*(*i′*) = *λ*(*i′*)*/p*(*i′*) and solving for *p*(*j′|i′*),

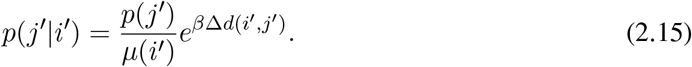

Enforcing the constraint that ∑_*j*∈*Ŝ*_ *p*(*j*|*i′*) = 1 and simplifying notation, we have

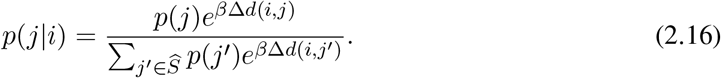

Before moving on, it is worth noting that Δ*d*(*i, j*) has a simple and intuitive interpretation. It is the difference in redundancy for component *j* when *i* is included versus when it is excluded, weighted by the relative importance of *i*.

Note that *p*(*j*) and *p*(*A|j*) depend on the choice of *p*(*j|i*). The final algorithm,

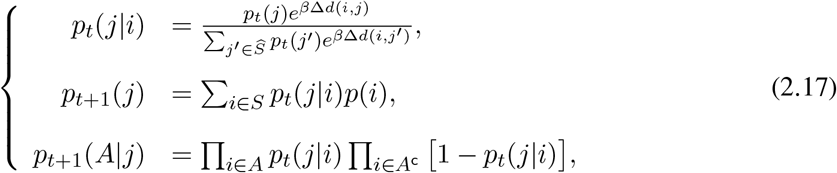

follows a similar alternating minimization scheme to the one developed by Blahut and Arimoto and generalized by Csiszár & Tsunády (1984), albeit with only local optimality guarantees similar to Tishby et al. (1999); Banerjee et al. (2005). See Appendix A and Alg. A1 for a complete derivation and description of the algorithm. The two practical issues with the algorithm are (1) the 2^*n*^ scaling of the number of subsets of *S* as *n* (the number of elements of *S*) increases, and (2) the general difficulty of estimating the mutual information between variables, let alone among multiple variables.

For the first issue, it is worth noting that there are non-trivial collective systems of empirical interest even for small *n*. Current computational hardware may permit exact computation up to around *n* ≈ 15 even on consumer hardware, which would be relevant for many experimental systems (as in e.g. Miller & Gerlai, 2007; Katz et al., 2011; Jolles et al., 2018). For larger systems, Monte-Carlo estimation of Δ*d*(*i, j*) can be readily employed, e.g. for *K* samples,

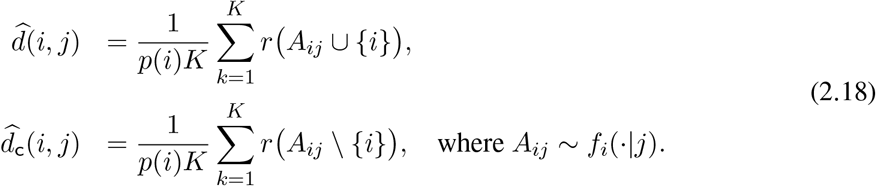

For large systems in particular initializing near good solutions may be helpful. In many systems we may expect elements to be spatially or temporally dependent, and use that prior knowledge to initialize reasonable clusters. However the preliminary results given in the next section do not employ any such strategy; we simply run the algorithm many times beginning with many different initial conditions and select the best solution generated.

While there is no exact universal solution to the practical difficulties of the second issue, we can proceed by maximizing a lower bound on component redundancy. For continuous random variables that are marginally Gaussian, the Gaussian mutual information is a lower bound on the total mutual information (Foster & Grassberger, 2011; Kraskov et al., 2004). Thus we can use

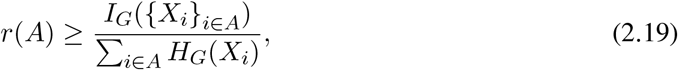

which is simple to compute in practice. When the random variables comprising the system are not marginally Gaussian, we can still use this bound by substituting copula transformed variables *G_i_* for *X_i_*, for which we enforce that each *G_i_* is Gaussian distributed. We emphasize that this transformation is valid and unique for any set of continuous random variables; this is guaranteed by Sklar’s theorem (Sklar, 1959), which ensures that the lower bound on redundancy given by Eq. 2.19 is applicable in general. The preliminary results in the next section are based on maximizing this Gaussian bound on redundancy using a copula transform to enforce Gaussian marginals.

### 2.3 Simulation experiments

We tested the proposed algorithm on two sets of data: simulations of schooling groups, and empirical data collected from the movements of schooling fish in a lab environment. The former allow us to control the dependency structure of the system, while the latter allows us to demonstrate applicability to empirical systems. Simulations used a simple model of coordinated movement based on attraction, alignment, and repulsion social forces (based on Romanczuk et al., 2012; Romanczuk & Schimansky-Geier, 2012; a description of the model and additional information on the simulation conditions can be found in Appendix B). Position and velocity data for independent groups of size *n* = 5, 10, and 20 were generated for a high (*η* = 0.2) and low (*η* = 0.15) noise conditions.

### 2.4 Empirical experiments

Movement data of fish comes from videos originally recorded by Katz et al. (2011). In that work, groups of 10, 30, and 70 golden shiners (*Notemigonus crysoleucas*) were purchased from Anderson Farms (www.andersonminnows.com) and filmed in a 1.2 × 2.1 m tank with an overhead camera. Videos were then corrected for lens distortion and fish were tracked using the same custom in-house software developed by Haishan Wu and used in Rosenthal et al. (2015). The software begins by detecting all individuals in each frame, then linking individuals across frames to form tracks. All tracks were manually corrected to ensure accuracy. Individual positions and velocities were estimated from these tracks using a 3^rd^ order Savitzky-Golay filter (Savitzky & Golay, 1964; similar to e.g. Harpaz et al., 2017) with a 7 frame smoothing window (videos were recorded at 30 fps). Interactions between fish are time-dependent: in results presented we chose a fixed window of ±15 s surrounding a given time *t* to estimate the dependency structure of the group.

## 3 Results

The algorithm outlined in the previous section requires specifying a parameter, *β*, which controls the relative importance of maximizing the average redundancy of the components as opposed to maximally compressing the original set of system elements. While it will be interesting to investigate the ‘soft-partitioning’ aspect of this approach in future work, here we simply consider the hard assignment case, which requires only that is *β* large. Fig. 2 (*Right*) illustrates this point, showing the stabilization of average component redundancy for *β* > 5. We found that *β* = 200 was sufficient to recover hard assignments in all cases tested here.^7^ Since relative redundancy ranges between 0 and 1 for any dataset, these parameter values should generalize well to other systems, and leaves the method free of parameter fine-tuning.

**Figure 2:**
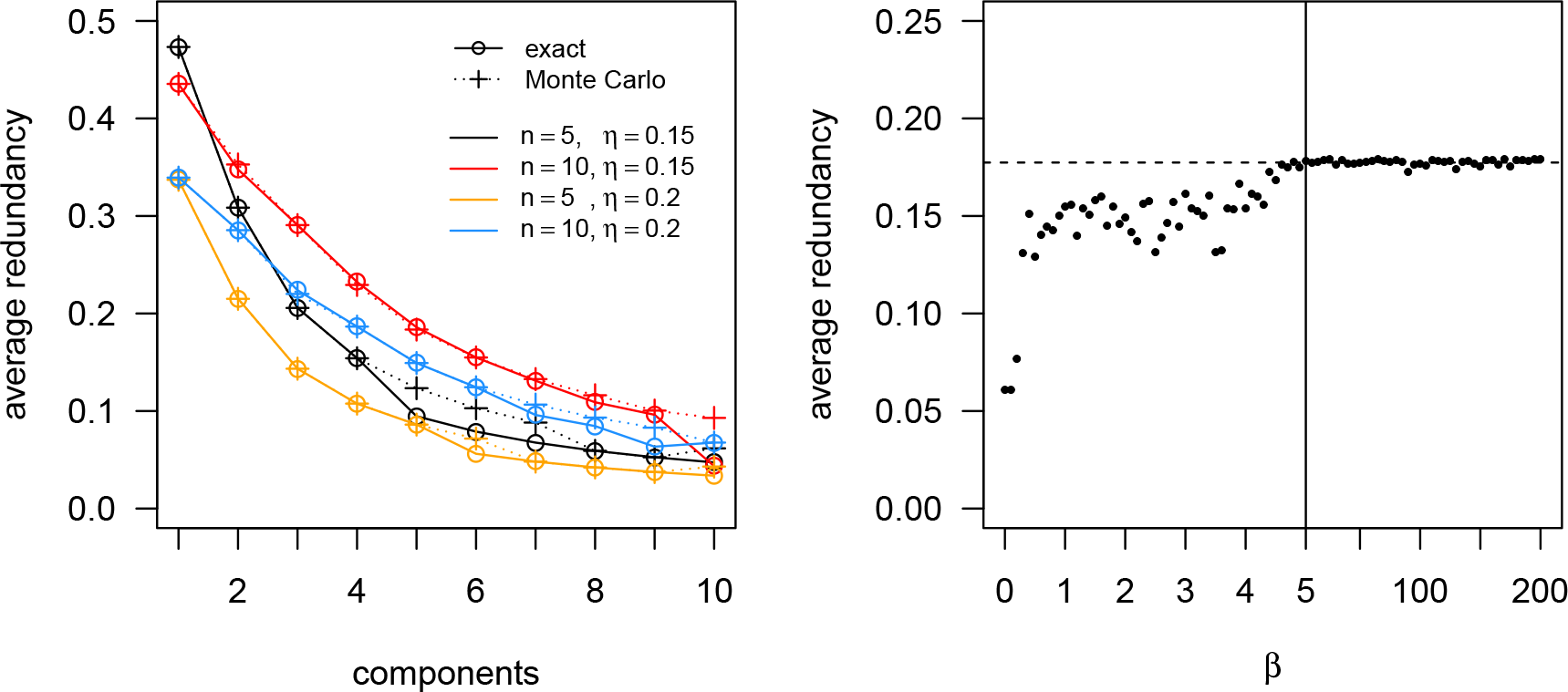
Algorithm implementation and parameter sensitivity. (*Left*) Comparison of exact and Monte Carlo estimates of Δ*d*(*i, j*), for single groups of size 5 and 10, for low and high noise conditions. (*Right*) Impact of the choice of on the *β* average redundancy of the recovered components for a single simulated group of size 10, high noise condition, searching for 5 components. Dotted line shows the mean of the solutions for *β* > 5.

To validate that the Monte-Carlo estimate of Δ*d*(*i, j*) employed is effective, we compared its behavior to exact computations of Δ*d*(*i, j*) for small system sizes (simulated groups of size 5 and 10). We ran each version of the algorithm for up to 10 components and took the best (maximum) average component redundancy achieved over 100 random initializations of the assignment matrix *p*(*j|i*). Fig. 2 (*Left*) shows that the results are in good agreement, and where there are discrepancies they tend to favor the Monte Carlo method.

We tested the Monte Carlo algorithm on simulated data in which the dependency structure of the simulated groups was known. For each test, we computed the best average component redundancy recovered for up to 10 components, again using 100 random initializations of the assignment matrix for each computation. Average component redundancies for single groups of size 5, 10, and 20 (Fig. 3 *Left*) have a peak at a single component, which includes all the elements of the system. Redundancies for two non-interacting groups, in pairs of matched size groups of 5, 10, and 20, have peaks at 2 components for size 10 and 20, with a plateau or slight decline for the pair of size 5 (Fig. 3 *Center*). Finally, the redundancies for a system of three non-interacting groups of mixed sizes 5, 10, and 20 was computed, with peaks at 3 and 4 components for high and low noise conditions, respectively (Fig. 3 *Right*).^8^ Taken together, this is evidence that the peaks and plateaus of the average component redundancies recovered by the algorithm do indeed reflect the dependency structure of the underlying system. It suggests that these features may be useful in identifying relevant structure in other systems, even those with less extreme dependency structures.

**Figure 3:**
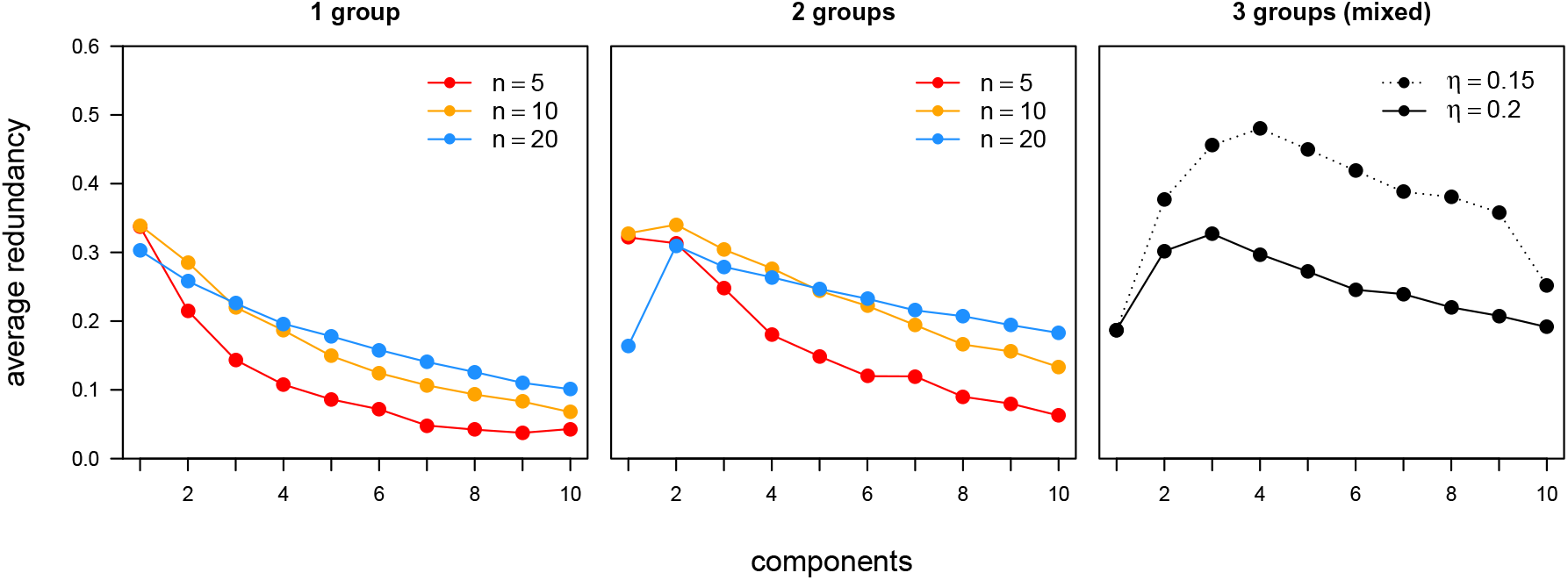
Partitioning results for simulations of 1, 2, or 3 independent (non-interacting) groups. (*Left*) For single cohesive groups of size (*n*) 5, 10, or 20, the average redundancy (y-axis) of the components decreases approximately monotonically as we increase the number of components. (*Center*) For two non-interacting groups of the same size, the average redundancy peaks, or at least plateaus, at two components. (*Right*) A mixed collection of three non-interacting groups, with sizes 5, 10, and 20, achieve peak average redundancy with three or four components, depending on the noise (*η*) used in the simulation. For comparison, the left two plots show results for *η* = 0.2 (the ‘high’ noise).

Fig. 4 illustrates the iterative generation of assignments for the algorithm in the mixed three group (high noise) case. Assignments change and harden until they converge on a (local) maximal average redundancy partition of the systems elements (*Left*). The assignments generated by the algorithm of system elements to components corresponds one-to-one with the original, non-interacting set of three groups (of sizes 5, 10, and 20) comprising the whole system (of total size 35). Positions of the elements of the system and their velocity vectors are shown for one time point, colored by the component they were assigned to (which corresponds to their original group), in Fig. 4 (*Left*). Note that, while the snapshot shown in Fig. 4 was chosen to show the three distinct groups, at many points in the simulation the positions, velocities, or both, overlapped between the three groups.

**Figure 4:**
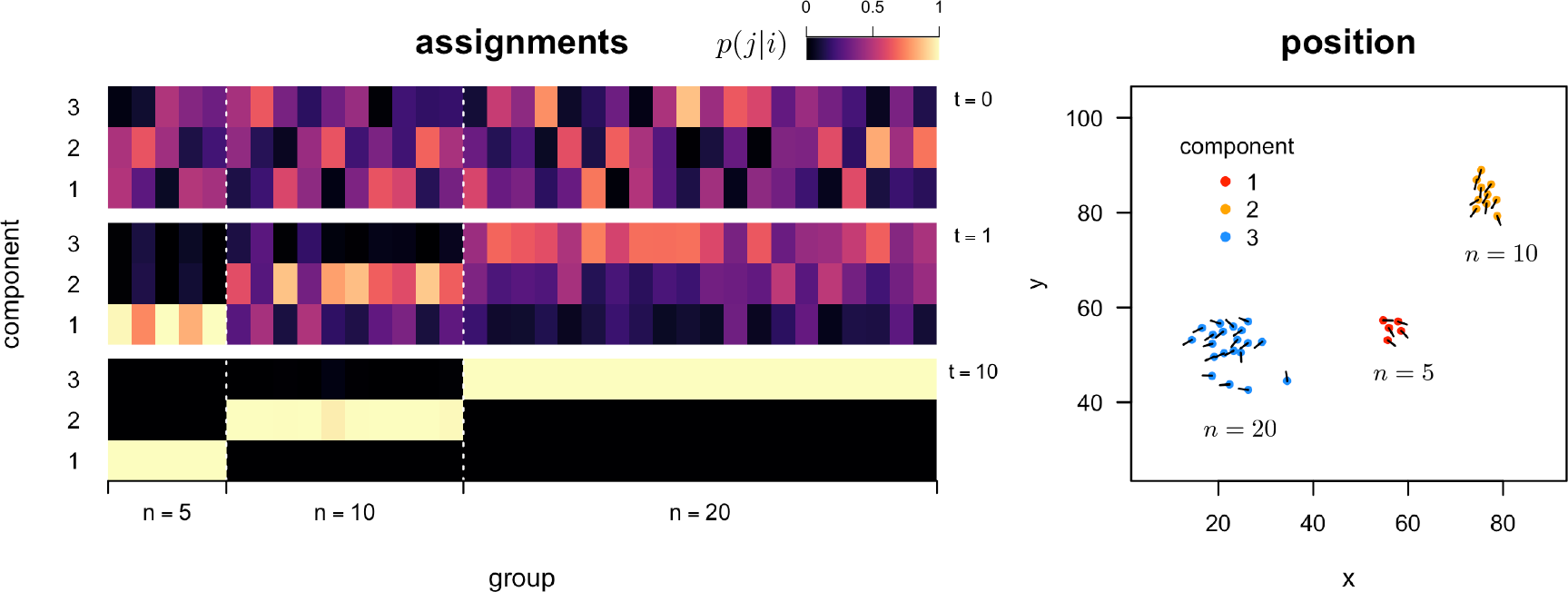
Generation of assignments for a mixture of three non-interacting simulated groups. (*Left*) Assignments generated by the proposed sequential algorithm for three components after initialization (*t* = 0), 1 iteration (*t* = 1), and 10 iterations (*t* = 10), at *top*, *middle*, and *bottom*, respectively. The color scale indicates the probability of assigning a member of a group (column) to a particular component (row), where low to high probability is coded dark to light (color scale top right). Original groupings of the system into its three non-interacting subsets are indicated on the x-axis. (*Right*) Two-dimensional positions (arbitrary units) of simulated system at one time point, color-coded by final component assignment; velocity vectors indicated by line segments.

Finally, we applied the algorithm to empirical data collected on fish schools to validate that the method is able to recover sensible results for strongly interacting groups and from non-simulated data. Fig. 5 shows that for fish, groups of size 10 interact strongly enough in at least one instance to be considered one coherent unit, while groups of size 30 are already large enough to have subsets that more strongly interact with one another than the rest of the group. Fish systems of size 70 do not always have a clear peak, but a broad plateau of possible subdivisions. The component assignments, positions, and velocities for a group of 30 fish is shown in Fig. 5 (*Right*) for three superimposed time points. Two of the components (shown as red and blue) show particularly coherent structure over the course of the 20 seconds of movement shown. Further work is needed to investigate the duration of substructure in fish schools, as well as the emergence and disappearance of components over time.

**Figure 5:**
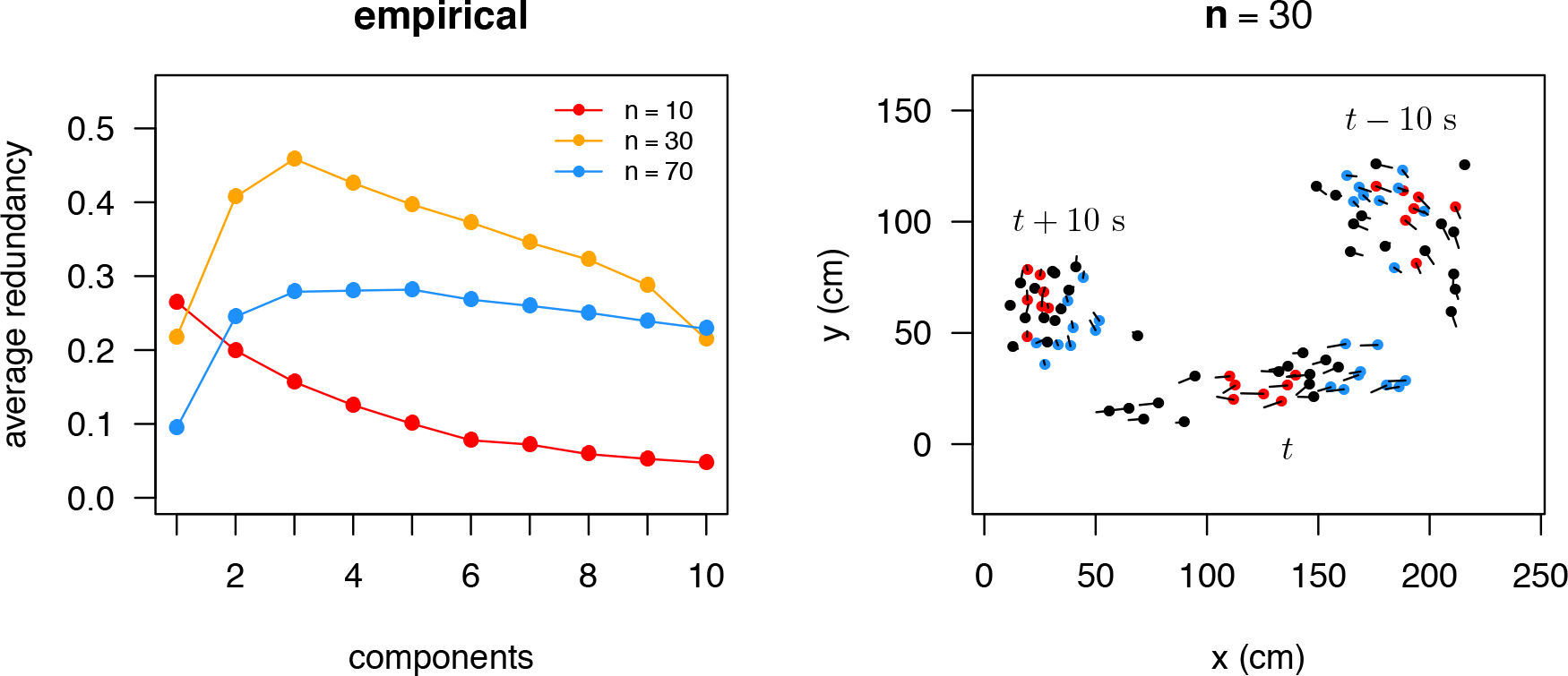
Redundant substructure for empirical fish schools. (*Left*) Average component redundancy as a function of the number of components, for fish groups of size 10, 30, and 70. (*Right*) Example partitioning of a group of size 30 fish into three components, colored black, red, and blue. Dots indicate the positions of the fish in a large (1.2 m × 2.1 m) arena, while line segments indicate the velocity vector of each individual. Positions (cm) and velocities are shown superimposed for the group at times *t* and *t ±* 10 *s*.

## 4 Discussion

There are a wide range of both general purpose clustering algorithms (see Jain, 2010; Xu & Tian, 2015) and network community detection methods (see Forunato, 2010), owing to a diversity of plausible clustering and community detection criteria. The justification for the average relative redundancy criterion presented here stems from its principled approach to the specific problem of quantifying collectivity, as argued at the beginning of this paper, and its demonstrated ability to identify the dependent structure of collective systems, as shown by the results obtained in the previous section. Usage of this criterion requires sufficient information for estimating the redundancy of any subset of the system, thus it cannot be used as a drop-in replacement for other clustering or community detection methods that operate on arbitrary similarity or correlation matrices. If the lower bound on redundancy given by Eq. 2.19 is to be used, then marginal normality must be enforced via copula transform or directly by the process generating the data.

In addition to its theoretical and empirical justification, the method is also computationally efficient, making it usable in practice. The proposed Monte-Carlo algorithm improves on the computational complexity of both the brute-force (check every partition) and naïve exact (sum over every subset) solutions, while achieving comparable results. It is instead limited by the cost of computing, for each element, log determinants for the Gaussian average redundancy bound. This reduces to matrix multiplication and thus scales (depending on the method) as *O* (*n*^3+1^*m*) or *O* (*n*^2.373+1^*m*), where *m* is the number of components, assuming a fixed number of Monte-Carlo samples and total iterations of the algorithm. In fact, this worst-case bound will almost never be achieved in practice, since it requires probabilistically sampling at least one assignment of all *n* elements to each of the *m* components when evaluating Δ*d*(*i,j*). The probability this occurs decreases exponentially in *n*.

There are many open questions left for future work. First, the identification of a peak in the average redundancy plot as a function of the number of components is only a heuristic. In some cases, as in e.g. the group of 70 fish studied here, there may be no obvious peak, and thus there may be more than one useful decomposition of the group. In other cases, depending on the question being asked, it may be more appropriate to divide the group into a given number of components regardless of the existence or position of a peak. Further theoretical work is needed on the significance of peaks or plateaus in the average redundancy plot; we present only empirical evidence of their utility here. Second, an investigation of these features as a function of the time-window chosen for computing the dependency structure may be important for understanding how the dependency structure of the group scales with time. It might be expected that this in fact plays a very important role, in that on short time-scales for many systems only very local interactions will matter, while at long enough time scales the system is best represented as only one component.

It may also be important to investigate the algorithm presented here in the context of generating a soft-partitioning of a system’s elements into partially overlapping components. Using intermediate values of *β* may allow the algorithm to find better average redundancy solutions ‘in-between’ *m* and *m* + 1 components, in which the assignments for some elements are split between some set of components. At the same time, since optimal sets of components are not guaranteed to be unique, it may be important to explore the set of equally (or nearly equally) optimal solutions as an ensemble of equivalent descriptions of a system. Moreover, exploring the range of solutions as the number of components varies may reveal whether or not the system exhibits some form of hierarchical dependency structure. In hierarchical systems we would expect components to be successively subdivided as the number of components increases, while in non-hierarchical systems this would not be the case.

One of the most interesting potential applications of the method may be to long time-series data for collective systems, in which the dependency structure of the group changes over time. Characterizing the natural decomposition of a system as a function of time may reveal important time-dependent mesoscopic features. How do the natural number of components of a system fluctuate in time, and how long do components persist? How do the components of a system interact as a function of time? These questions are central to the study of collective systems and can now be addressed quantitatively via the method introduced here.

## A Algorithm

Here we give an expanded account of the redundancy compression algorithm.

### A.1 Rate-Distortion Compression

Classical rate-distortion theory treats the following optimization problem:

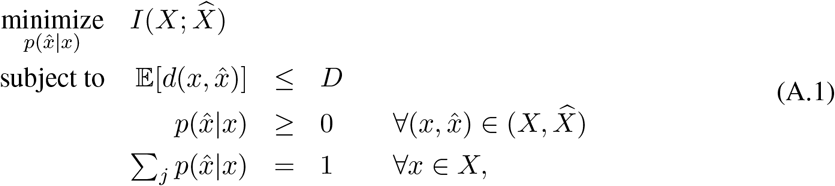

where

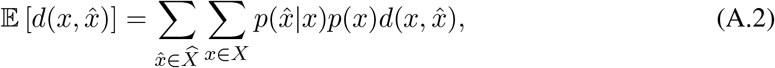

and *p*(*x*) is given. The problem as stated is not convex due to the form of *I*(*X*; *X̂*). However, writing the objective as

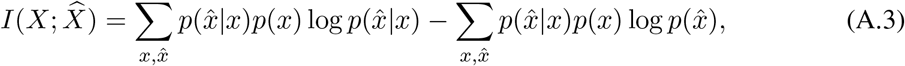

it is clear that the problem is convex when varying *p*(*x̂|x*) or *p*(*x̂*) separately, holding the other constant. Since the distortion constraint, 𝔼[*d*(*x*, *x̂*)] is convex in *p*(*x̂|x*), the problem can be restated as a convex double minimization of the form

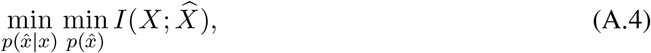

which is minimized for fixed *p*(*x̂|x*) by

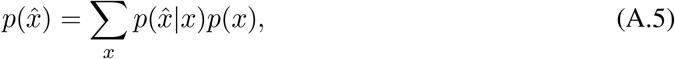

and for fixed *p*(*x̂*) by

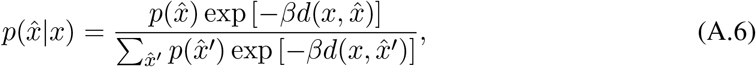

(see Blahut, 1972; Arimoto, 1972; Cover & Thomas, 2006). This leads to the classic Blahut-Arimoto algorithm, which, by iterative application of these two self-consistent equations for a given *β*, converges to an optimal solution point on the rate-distortion curve with tangent slope equal to *β*.

### A.2 Redundancy Compression

In this paper, we are interested in a similar problem:

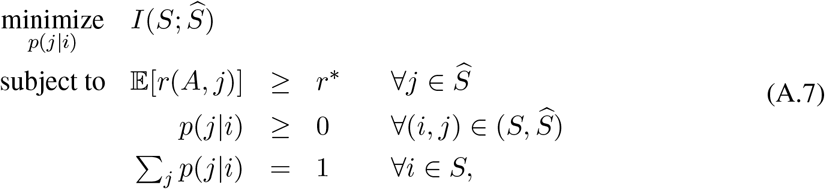

where

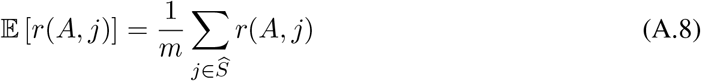

and

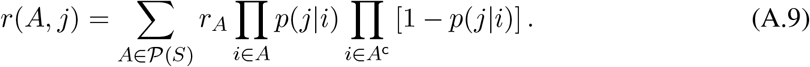

The fixed 1/*m* weighting of the marginal importance of each component, *j*, in the redundancy constraint, 𝔼[*r*(*A, j*)], is a minor variation from the classical rate-distortion problem. The important difference is that the *r*(*A, j*) inequality constraint is not convex with respect to *p*(*j|i*). However, with change of variables *b_A_* = log *r_A_*, *y_ij_* = log *p*(*j|i*), and *ȳ_ij_* = log [1 − *p*(*j|i*)], we can define

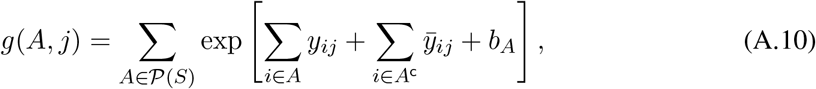

where *r*(*A, j*) = *g*(*A, j*), with *g*(*A, j*) convex with respect to *y_ij_* and *ȳ_ij_* and invariant with respect to *p*(*j|i*) or *p*(*j*).

This gives the equivalent minimization problem:

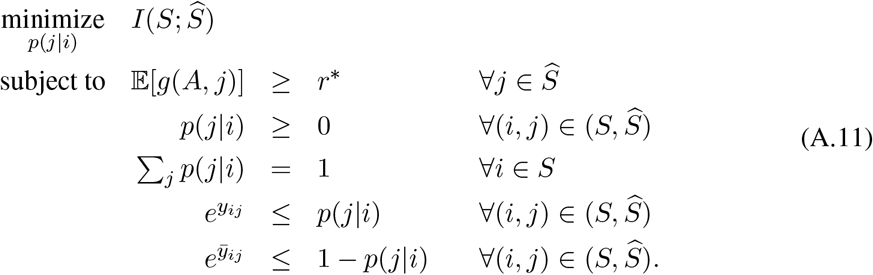

Setting aside non-negativity constraints on *p*(*j|i*) (these will be enforced by the form of the solution), we have the functional

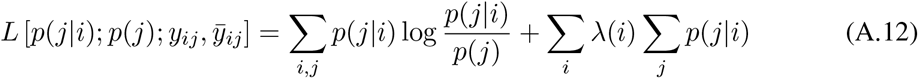

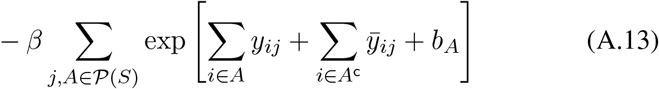

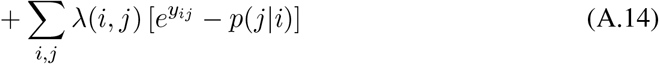

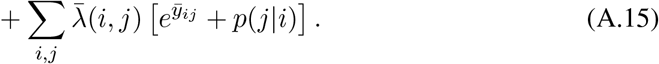

We can then restate the original non-convex problem in terms of two convex minimizations and one quasiconvex minimization,

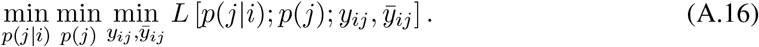

Note that, similar to Tishby et al. (1999), the problem is not jointly convex and thus there is no guarantee of a unique global solution as in the rate-distortion case. Nevertheless, the marginal (quasi-)convexity admits an efficient iterative algorithm for identifying (locally) optimal solutions, similar to Tishby et al. (1999).

Taking the derivative of *L* with respect to *p*(*j|i*) and setting equal to zero, we arrive at

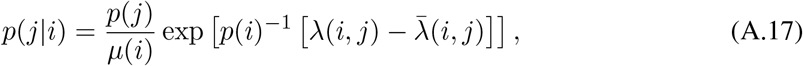

where *µ*(*i*) just normalizes the distribution over *j* for a given *i*. Taking the derivative of *L* with respect to *y_ij_* and setting equal to zero, we have

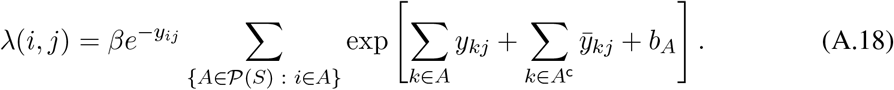

Doing the same for *ȳ_ij_* gives

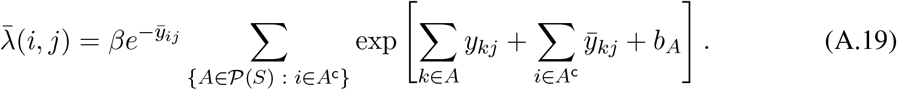

Subtracting the two equations, we have

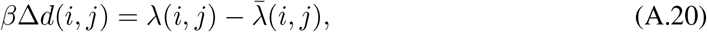

which is equivalent to the definition of *d*(*i, j*) in the main text. Substituting into Eq. A.17 produces

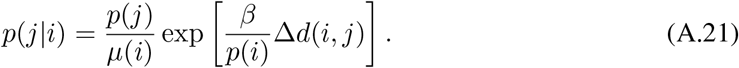

This gives the minimizing values of *L* with respect to *p*(*j|i*) for fixed *p*(*j*), *y_ij_*, and *ȳ_ij_*, as in Blahut (1972); Arimoto (1972); Tishby et al. (1999); Banerjee et al. (2005). The minimizing values of *L* with respect to *p*(*j*) are the same as in classical rate-distortion theory and are given by

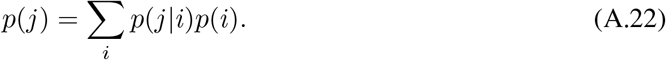

The minimizing value of *L* with respect to *y_ij_* and *ȳ_ij_* under the constraints that *e^yij^* ≤ *p*(*j|i*), and *e*^*ȳij*^ ≤ [1 *− p*(*j|i*)], is simply

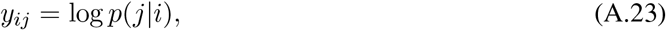

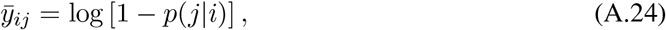

since the monotonically decreasing A.13 will achieve its minimum for the least negative values of *y_ij_* and *ȳ_ij_*, which puts them up against their constraints.

### A.3 Generalization

It is clear from the form of *g*(*A, j*) that the only requirement of the measured property, *b_A_*, of any set, *A* ∈ *S*, is that it is non-negative. Thus this same method may be employed for measures on sets other than redundancy, in the same way that rate-distortion theory treats generic measures of distortion. On the other hand, when the measured property offers certain kinds of additional structure, as in e.g. the case of an average similarity (Slonim et al., 2005) measure, then other efficient solutions may be possible.

One variant to the sequential update of *p*(*j|i*) as listed in Alg. A1 is to modify every *p*(*j|i*) in parallel, which may be advantageous for some multiprocessor configurations. In practice, for convergence with simultaneous updating it appears to be important to introduce a slowdown factor, *α*, to control the update of *p_t_*(*j|i*), i.e. using

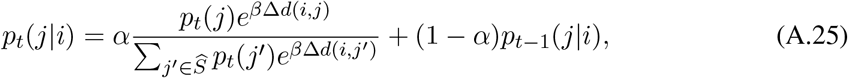

where *t* is the current iteration of the algorithm. The slowdown operates in a manner analogous to the learning rate in gradient descent optimization problems.

**Algorithm A1.**
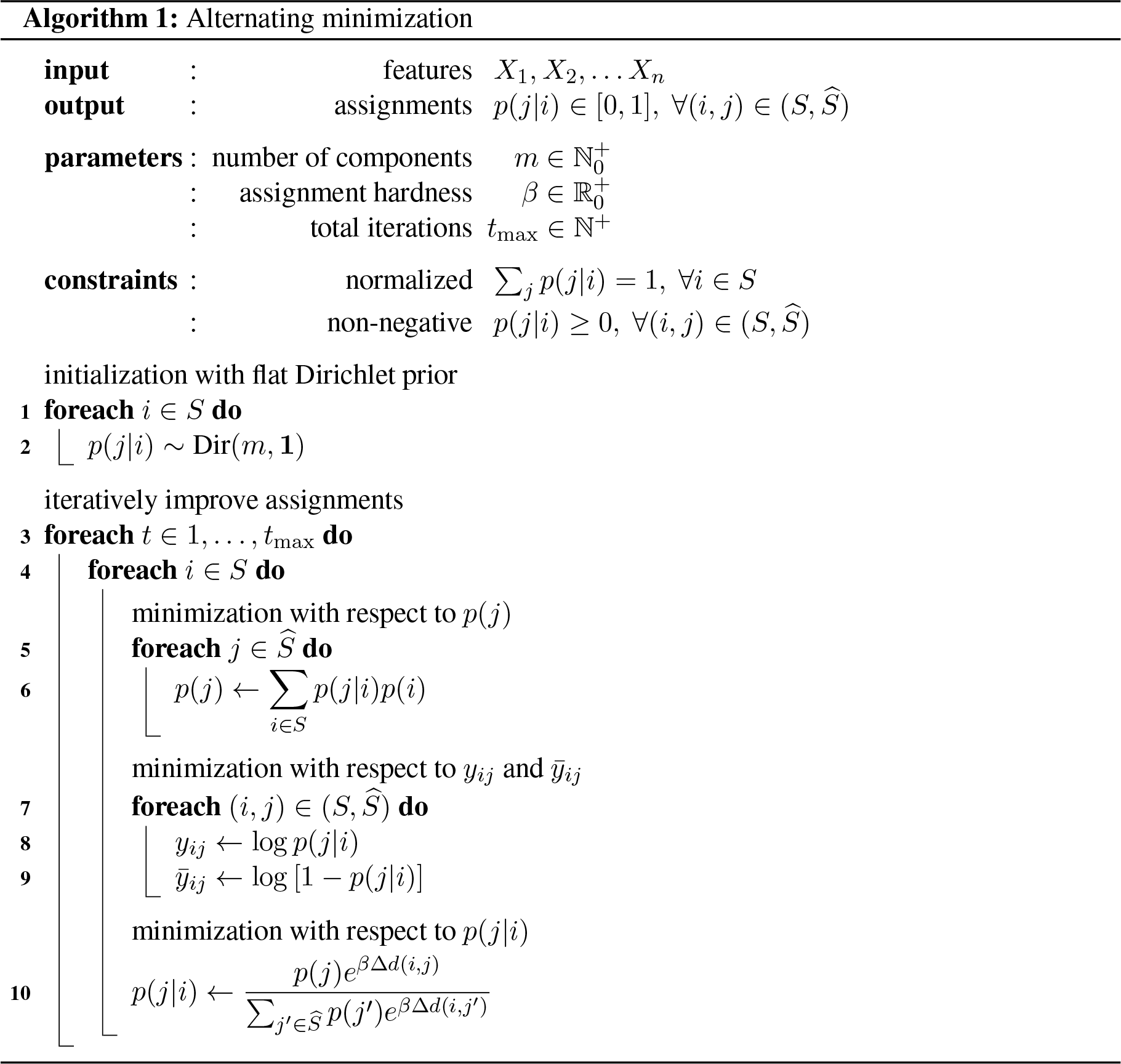
ℕ^+^ are the positive integers, while ℕ^+^_0_, ℝ^+^_0_, are the non-negative (positive including zero) integers and real numbers, respectively. For hard clustering, *β* just needs to be large. Parameter *t*_max_ needs to be large enough for convergence; alternatively, it can be replaced by a criterion based on a minimum difference in improvement between iterations. Lines 2 and 10 are to be understood as vector operations over the set *j* ∈ *Ŝ*

Like *β, α* does not require fine-tuning. It just needs to be small enough to allow for convergence, without being too small so as to allow the algorithm to converge in a reasonable number of iterations. While a more systematic investigation may be useful in identifying an efficient *α*, we found that *α* = 0.1 and *t* = 200 iterations was sufficient to ensure convergence for all the numerical results presented in the main text. In many cases a stable assignment is reached much earlier than after 200 iterations, and in general a stopping criteria based on the difference between assignments from one iteration to the next could be employed, though we did not do so here.

## B Simulation

The agent-based model used in this paper for generating schooling motion with known dependency structure is based on the three-zone-model introduced by Couzin et al. (2002). Each agent moves at a constant speed *s*_0_ and responds to its conspecifics by changing its direction of motion. The interactions between individuals are governed by three basic social forces: long-range attraction, short-range repulsion, and intermediate-range alignment. However, there are two main differences from the original Couzin model: (1) the model is formulated in terms of stochastic differential equations with effective social forces (see Romanczuk et al., 2012; Romanczuk & Schimansky-Geier, 2012); and (2) instead of discrete zones, we use overlapping social forces, whereby repulsion dominates at short distances (*r_ij_* < *r*_rep_), attraction dominates at long distances *r_ij_* < *r*_att_, and the alignment contribution overlaps with attraction and repulsion up to intermediate ranges (*r_ij_* < *r*_alg_), whereby *r*_rep_ < *r*_alg_ < *r*_att_.

### B.1 Model formulation

We simulate the movement of a group of *n* agents via a set of 2*n* (stochastic) differential equations. The agents move in a quadratic domain of size *L × L* with periodic boundary conditions. The dynamics of each agent (in 2d) are described by the following equations of motion (*i* = 1, …, *n*):

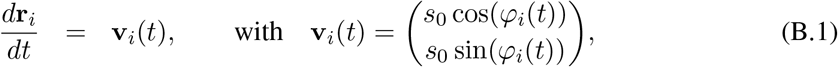

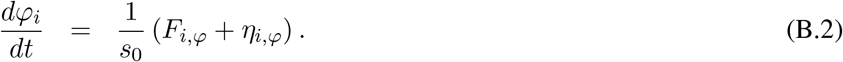

Here **r**_*i*_, and v_*i*_ are the Cartesian position and velocity vectors of each agent, with *s*_0_ being the (constant) speed of agent *i*. Furthermore, *η_i,φ_* are Gaussian white noise terms accounting for randomness in the turning motion of individuals, and **f**_*i,φ*_ are the projections of the total social forces inducing turning behavior, where

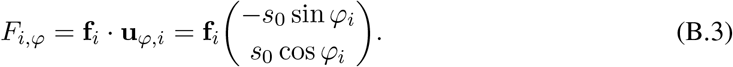

The total effective social force is a sum of three components, **f**_*i*_ = **f**_*i*,rep_ + **f**_*i*,alg_ + **f**_*i*,att_,

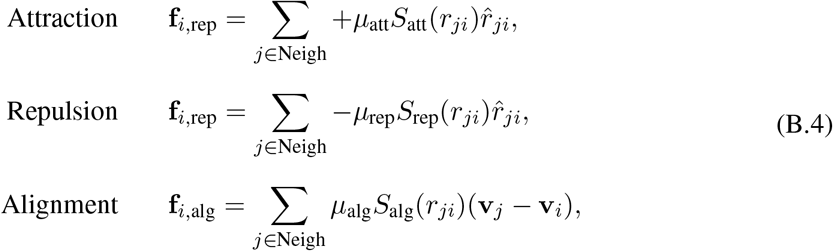

with *r̂* = **r***/|r|*. The strength of the different interactions is set by a constant *µ_X_* and a sigmoid function of distance, which goes from 1 to 0, with the transition point at *r_X_* and steepness *a_X_*:

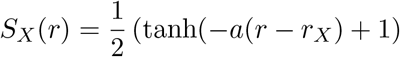

**Algorithm B1.**
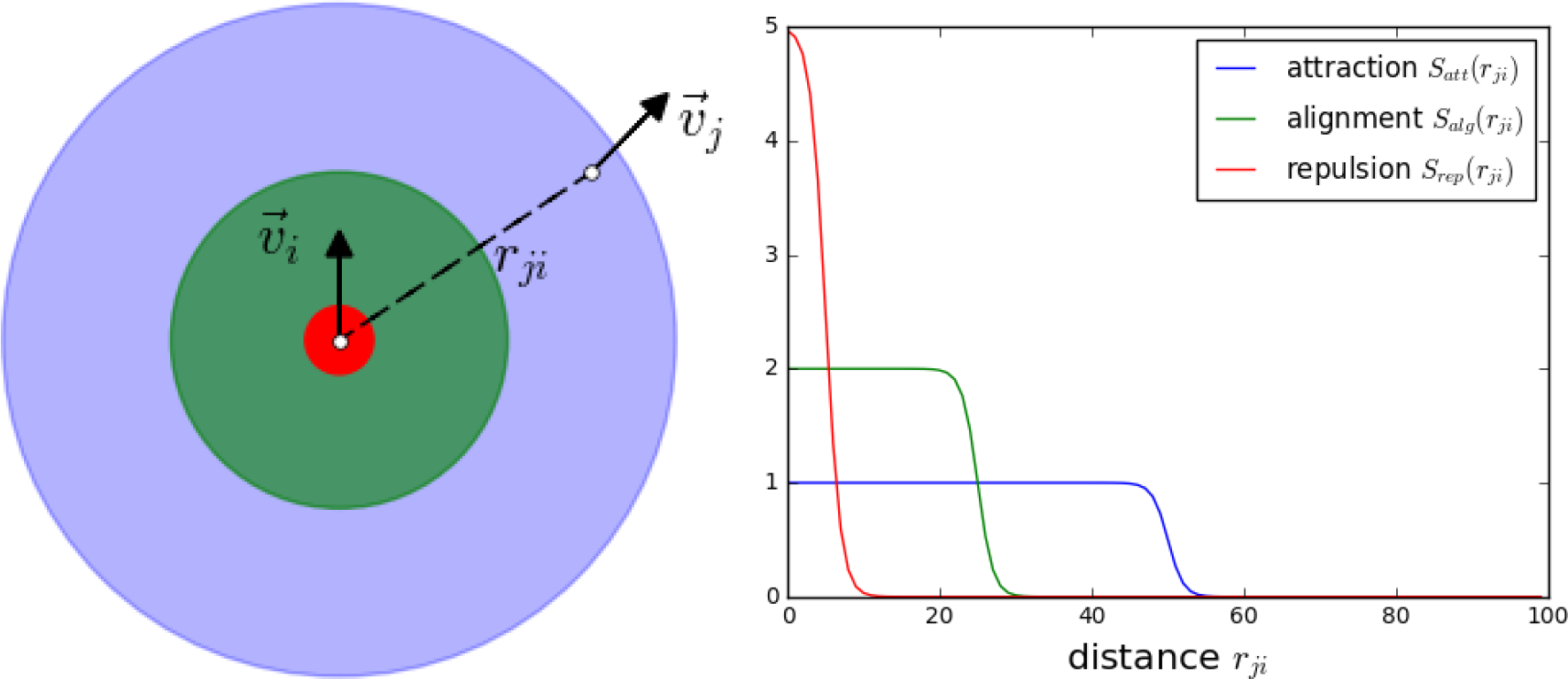
(*Left*) Schematic of the effective social interactions, with repulsion dominating at short distances (red zone), attraction dominating at large distances (green zone) and main contribution of alignment at intermediate ranges (blue zone). (*Right*) The strength of the different social forces versus distance for the different interactions.

(see Fig B1).

The stochastic differential equations for the direction of motion of individual agents are solved by a simple Euler-Maruyama method:

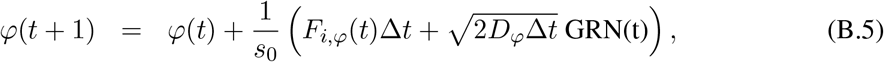

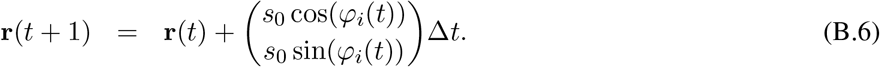

### B.2 Numerical experiments

We simulated independent groups of three different sizes, *n* = 5, 10, and 15, wherein it was possible for each agent to interact with the distance dependent effective forces with all other agents within the group. The initial conditions were always a random distribution of agents in the simulation domain with random initial direction of motion. In order to ensure formation of a single cohesive group we set the attraction range to be larger then the domain size *r*_att_ *> L*. In all simulation runs considered here, we obtained for the used parameters (see Tab. 1) a single polarized group after a transient time of *t* < 400. Thus for our analyses we used only data for *t* > 400.

**Table 1.** Parameter values used in the simulations.

| Parameter | Symbol | Value |
| --- | --- | --- |
| domain size | $L$ | 100 |
| repulsion range | $r_{\text{rep}}$ | 1.0 |
| attraction range | $r_{\text{att}}$ | 100.0 |
| alignment range | $r_{\text{alg}}$ | 5.0 |
| repulsion strength | $\mu_{\text{rep}}$ | 2.0 |
| attraction strength | $\mu_{\text{att}}$ | 0.3 |
| alignment strength | $\mu_{\text{alg}}$ | 0.8 |
| steepness of interaction function | $a$ | 10 |
| speed of individuals | $s_0$ | 1.0 |

## Footnotes

1 As noted by Watanabe (1960), its significance as a potential measure of organization stretches back still further, to at least Rothstein (1952).

2 Meaning that they share the same outcome.

3 If *m > n* then some components will necessarily be empty.

4 The use of *i* and *j* as elements of *S* and *Ŝ*, respectively, will follow this convention in the rest of the paper.

5 Unless stated otherwise, the complement of a set is taken with respect to *S*, i.e. *A*^c^ = {*k* ∈ *S*: *k* ∉ *A*}.

6 A consequence of the differences with the standard rate-distortion formulation is that we should not expect *R*(*r\**) to necessarily behave similarly to *R*(*D*) as we vary *r\** and *D*, respectively.

7 Using the simultaneous updating variant of the algorithm, see Appendix A.

8 In both noise conditions all three non-interacting groups were split into separate components. In the low noise condition, the group of 20 was further subdivided into two components.

